# The Long and the Short of Serotonergic Stimulation: Optogenetic activation of dorsal raphe serotonergic neurons changes the learning rate for rewards

**DOI:** 10.1101/215400

**Authors:** Kiyohito Iigaya, Madalena S. Fonseca, Masayoshi Murakami, Zachary F. Mainen, Peter Dayan

**Affiliations:** Gatsby Computational Neuroscience Unit, UCL, London W1T 4JG, UK; Max Planck UCL Centre for Computational Psychiatry and Ageing Research, London WC1B 5EH, UK; Division of Humanities and Social Sciences, California Institute of Technology, Pasadena, CA 91125; Champalimaud Neuroscience Programme, Champalimaud Centre for the Unknown, Avenida de Braslia, 1400-038 Lisbon, Portugal

## Abstract

Serotonin plays an influential, but computationally obscure, modulatory role in many aspects of normal and dysfunctional learning and cognition. Here, we studied the impact of optogenetic stimulation of dorsal raphe serotonin neurons in mice performing a non-stationary, reward-driven, foraging task. We report that activation of serotonin neurons significantly boosted learning rates for choices following long inter-trial-intervals that were driven by the recent history of reinforcement.

## 1 Introduction

Learning from the outcomes of past actions is crucial for effective decision-making and thus ultimately for survival. In the case of important outcomes, such as rewards, ascending neuromodulatory systems have been implicated in aspects of this learning due to their pervasive effects on processing and plasticity. Of these systems, perhaps best understood is the involvement of phasically-fluctuating levels of dopamine activity and release in signalling temporal difference [55] prediction errors for appetitive outcomes [46, 50]. Since prediction errors are a key component of reinforcement learning (RL) algorithms, signalling mismatches between outcomes and predictions, this research has underpinned and inspired a large body of theory on the neural implementation of RL.

From the early days of investigations into aversive processing in Aplysia [27], serotonin (5-HT) has also been implicated in plasticity. This is broadly evident in the mammalian brain, from the restoration of the critical period for the visual system of rodents occasioned by local infusion of 5-HT [58] to the impairment of particular aspects of associative learning arising from 5-HT depletion in monkeys [7, 59]. Despite theoretical suggestions for an association with aversive learning [19, 53, 13, 6, 15], direct experimental tests into serotonin’s role in RL tasks have led to a complex pattern of results [12, 52, 42, 45, 22, 11]. For instance, recent optogenetic studies reporting that stimulating 5-HT neurons could lead to positive reinforcement [42] do not appear to be consistent with other optogenetic findings, which instead suggest an involvement with patience [22, 45] and even locomotion [11].

Here, we study a different aspect of the involvement of 5-HT in RL. Although prediction errors are necessary signals for learning, they are not sufficient. This is because there is flexibility in setting the learning rate, i.e., the amount by which an agent should update a prediction based on such errors. The learning rate determines the timescale (e.g. how many trials) over which reward histories are integrated to assess the value of taken actions. 5-HT can readily influence learning rates through its interaction with dopamine [21]; and indeed, there is evidence that animals adapt the timescales of plasticity to the prevailing circumstances [16, 4, 47, 62, 30], and also consider more than one timescale simultaneously [10, 38, 23, 31]. 5-HT could be involved in some, but not other, timescales. It could also be associated with some, but not other, of the many decision-making systems [14, 25, 9, 40] that are known to be involved in RL.

We therefore reanalyzed experiments in which mice performed a partially self-paced, dynamic foraging task for water rewards [22]. In this task, 5-HT neurons in the dorsal raphe nucleus (DRN) were optogenetically-activated during reward delivery in a trial-selective manner. The precise control of the timing and location of stimulation offered the potential of studying in detail the way in which 5-HT affects reward valuation and choice. We used methods of computational model comparison to examine these various possible influences. We first noted a substantial difference in the control of actions that followed short and long intertrial intervals: only the latter were influenced by extended reward histories, as expected for choices driven by conventional RL. We then found that the learning rate associated with these (latter) choices was significantly increased by 5-HT stimulation.

## 2 Results

### 2.1 Animals showed a wide distribution of inter-trial-intervals (ITIs)

We reanalyzed data from a dynamic foraging or probabilistic choice task in which subjects faced a two-armed bandit [22]. Full experimental methods are given in that publication. Briefly, the subjects were four adult transgenic mice expressing CRE recombinase under the serotonin transporter promoter (SERT-Cre) and four wild-type littermates (WT) [22]. In this task (**Figure 1a**), mice were required to poke the center port to initiate a trial. They were then free to choose between two side ports, where reward was delivered probabilistically at both ports on each trial (on a concurrent variable-ratio-with-hold schedule [39]). On a subset of trials, when mice entered a side port, one second of photo-stimulation was provided to DRN 5-HT neurons via an implanted optical fiber (**Figure 1b**). ChR2-YFP expression was histologically confirmed to be localized to the DRN in SERT-Cre mice (**Figure 1c**) [22].

**Figure 1:**
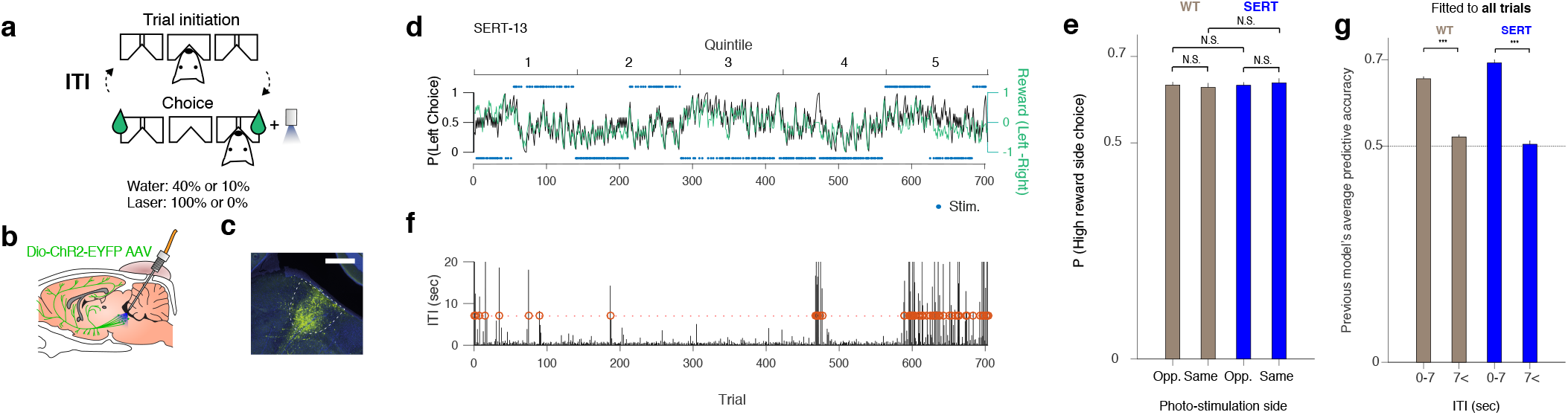
(**a**). Schematic diagram of trial events in the task. On each trial, a mouse was required to enter the center port (Trial initiation) and then move to one of the side ports (Choice). A reward might be delivered at the side port according to a variable-ratio-with hold-schedule. The next trial started when the mouse entered the center port. The inter-trial-interval (ITI) is defined as the time from when the mouse left the side port until it entered the center port to initiate the next trial. In a given block of trials, one side port was associated with a higher reward probability per trial (0.4) than the other (0.1); although following delivery, rewards were held (but not accumulated) until collected. Furthermore, during a block, photo-stimulation (12.5 Hz, 5 mW for 1 s) was always delivered as soon as the mouse entered just one of the side ports. (**b**). A schematic of the optogenetic stimulation. DRN neurons were infected with viral vector AAV2/1-Dio-ChR2-EYFP. In SERT-Cre mice, 5-HT neurons expressed ChR2-YFP (green) and could be photoactivated with blue light that was delivered by an optical fiber implant. (**c**) A fluorescence image of a parasagittal section shows localized ChR2-YFP expression (YFP = green, DAPI = blue) in the DRN. The white bar indicates the scale of 500 mm. (**d**). Time course of mouse choice behavior in an example session. The probability across trials of choosing the left port (black solid line) is overlaid with the collected reward bias (green line) for an example mouse, SERT-13. The choice probability and the reward bias were computed by a causal half Gaussian filter with a standard deviation of 2 trials. For the reward bias, a stream of values 1 (a reward from Left), −1 (a reward from Right), and 0 (no reward) was used for the estimation. The top (bottom) light blue dots indicate photo-stimulation at Left (Right) port. (**e**). The probability of choosing the higher water probability side is shown for the blocks in which the photo-stimulation was assigned to the opposite side from the higher water probability side (Opp.), and for the blocks in which the photo-stimulation was assigned to the same side (Same). The difference within WT mice, within SERT-Cre mice, and between WT and SERT-Cre mice for either condition were not significant. The error bars indicate the mean ± SEM over sessions. (**f**). Inter-trial-intervals (ITIs) in the same session as c. The red circle indicates trials with long ITIs (> 7 sec). (**g**). The average predictive accuracy of the existing reward- and choice-kernel model [39, 22] when fitted to all trials. This model captures a form of win-stay, lose-shift rule. Choices following short ITIs (≤ 7 s) were well predicted by the model, while choices following short ITIs (> 7 s) were not. The difference between short and long ITIs was significant for both WT and SERT mice [permutation test. *p <* 0.001 indicated by three stars.]. Data and images from [22]

Following previous experiments in macaque monkeys [54, 10, 39], the probability that a reward is associated with a side port per trial was fixed in a given block of trials (Left vs Right probabilities: 0.4 vs 0.1, or 0.1 vs 0.4). Once a reward had been associated with a side port, the reward remained available until collection (although multiple rewards did not accumulate). Photostimulation was always delivered at one of the side ports in a given block (Left vs Right probabilities: 1 vs 0, or 0 vs 1). Block changes occurred every 50-150 trials and were not signaled, meaning that animals needed to track the history of rewards in order to maximize rewards.

As previously reported [22], subject's choices tended to follow changes in reward contingencies (**Figure 1d**), exhibiting a form of matching behavior [54, 10, 39]. A deterministic form of matching behavior can maximize average rewards in this task [49, 43, 32, 31] because the probability of getting a reward increases on a side as the other side is exploited (due to the holding of rewards). For more behaviorally realizable policies, slow learning of reward contingencies has been shown to be beneficial to increase the chance of obtaining rewards [31].

We confirmed the results of previous analyses [22] showing that the optogenetic stimulation of DRN neurons did not appear to change the average preference of the side ports (**Figure 1e**). The animals’ preference for the side port that was associated with a higher water probability was not affected by the side which was photo-stimulated. However, these analyses do not fully take advantage of the experimental design in which photo-stimulation was delivered on a trial-by-trial basis. The latter should allow us to examine whether the effect of stimulation is more prominent on a specific subset of trials.

### 2.2 Fast or slow: ITI duration determined decision policy

The task contained a free operant component in that the subjects were free to initiate each trial. This resulted in a wide distribution of inter-trial-intervals (ITIs). It was notable that some ITIs were substantially larger than others (**Figure 1f**). To quantify this effect, we separated short from long ITI trials using a threshold of 7 seconds (we consider other thresholds below; values greater than 4 seconds led to equivalent results).

**Figure S1** reports the mean proportions of long ITI trials in WT and SERT-Cre mice. The frequency of long ITI trials was slightly, but statistically significantly, different between WT and SERT; however, this appears not to be due to stimulation itself, as control analysis showed that stimulation itself did not significantly change the ITI that followed (**Figure S2**). We also found that long ITI trials were most common in the last part of each experimental session, but were also seen in earlier parts of each session (**Figure S3**).

Previous studies have suggested a relationship between the duration of an ITI and the nature of the subsequent choice. For example, subjects have been reported to make more impulsive choices after shorter ITIs [48]. Another studies have shown that perceptual decisions are more strongly influenced by more venerable prior experience when working memory was disturbed during the task [1]. Here, we hypothesized that choices following short ITIs might also be more strongly influenced by the most recent choice outcome compared to those following long ITIs, since the outcome preceding a short ITI is more likely to be kept in working memory until the time of choice.

To investigate this, we first exploited an existing model of the behavior on this task [39, 22]. This is a variant of an RL model which separately integrates reward and choice history over past trials, subject to exponential decay [39]. This model captures a form of win-stay, lose-shift rule [3, 61] when time constants are small.

We found that choices following short ITIs (ITIs < 7 s) were well-predicted by this previously validated model (see Methods for details) (**Figure 1g**). Further, the time constants of the model were indeed very short (Reward kernel: 1.4 trials for WT, 1.9 trials for SERT-Cre mice; Choice kernel: 1.3 trials for WT, 1.2 trials for SERT-Cre mice). This suggests choices followed a form of win-stay, lose-shift rule [3, 61]. The difference of the reward time constant between WT and SERT-Cre mice was significant (p < 0.01, permutation test) but very small (< 1 trial), while the choice time constant was not. This paltry difference in reward time constant suggests a slightly smaller learning rate for the SERT-Cre mice, since the learning rate is inversely proportional to the time constant.

However, choices following long ITIs (ITIs > 7 s) were not well predicted by the same model (**Figure 1g**), suggesting that choices following short ITIs and long ITIs are qualitatively different. It also suggests that choices following long ITIs cannot be accounted for by a short-term-memory-based win-stay lose-switch strategy.

We hypothesized that choices following long ITIs might reflect slow learning of reward history over many trials [36, 31]. Indeed, by complexity-adjusted model comparison (integrated BIC) [29, 34], we found that choices following ITIs > 7 s were best described by a standard RL model (**Figure S4**). This analysis supported our hypothesis that choices following long ITIs are influenced by a relatively long period of reward history compared to choices following short ITIs. It is also worth noting that in contrast to the short ITI model, in which memory decays rapidly every trial regardless of choice, the standard RL model does not change the value of an option as long as the option is not selected. This suggests that different memory mechanisms may be involved in the decisions following short and long ITIs.

### 2.3 Optogenetic stimulation of DRN 5-HT neurons increased learning rate: Model-agnostic analysis

Given our original hypothesis that serotonin modulates the RL learning rate, we predicted that optogenetic stimulation of DRN 5-HT neurons would have a stronger impact on choices following long ITIs, since those choices appear to be more sensitive to learning over long trial sequences.

To test this, we first conducted the model agnostic analysis described schematically in **Figure 2a**. To assess how reward history with or without photo-stimulation affected choice following long ITIs, we estimated correlations between the temporal evolution of the reward bias and the choice bias for trials preceded by long ITIs. We did this separately for trials with and without serotonin photo-stimulation.

**Figure 2:**
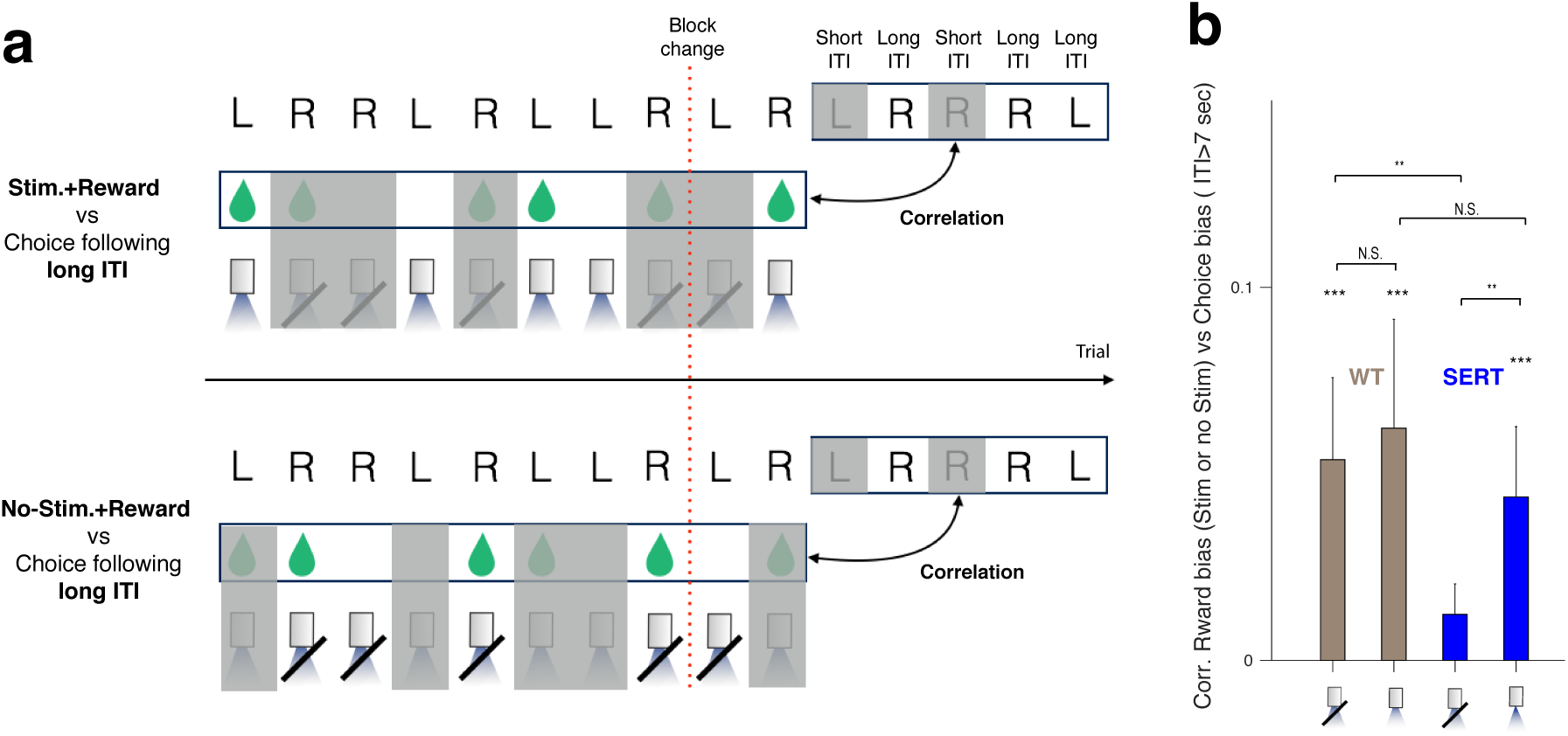
(**a**). Schematic diagram of model-agnostic analysis. The correlation between the choices following long ITIs (window = 5 trials) and the reward bias (window = 10 trials) was estimated using adjacent sliding windows. The reward bias was estimated on trials only with (top) or without (bottom) photo-stimulation. The windows were shifted one trial at a time. The greyed-out trials are the ones that are ignored for the assessments. Note that, due to the task design in which photo-stimulation is associated with only one side (Left or Right) in a given block, in some moving windows reward bias had to be computed from one side only. Thus we assigned +1 (respectively −1) to a reward from Left (Right) and no-reward from Right (Left) when we computed reward bias. We aware that this is not a perfect measure for reward bias; but we still expect finite correlations since reward rates from the Left choice and the Right choice are on average negatively correlated by the task design in a given block (reward probability: 0.1 vs 0.4). The correlation was estimated separately for each mouse. (**b**). Model-agnostic analysis suggests that the impact of reward history on choices following long ITIs was modulated by optogenetic stimulation. The x-axis indicates if the reward bias was computed over trials with or without photo-stimulations. The stars indicate how significantly the correlation is different from zero, or the correlations are different from each other, tested by a permutation test, where estimated reward bias was permuted within or between conditions. Three stars indicates *p <* 0.001. The error bars indicate the mean ± SEM of data.

As seen in **Figure 2b**, we found significant correlations between reward and choice bias for all conditions. Importantly, there was a significant effect of serotonin stimulation on the magnitude of the correlation. That is, for the SERT-Cre mice the correlation was larger for stimulated trials. This suggests that optogenetic stimulation of 5-HT neurons modulated learning about reward history, which in turn affected choices following long ITIs. The equivalent analysis for short ITIs (**Figure S10**) showed that they were not affected by serotonin stimulation in the same way. Indeed, a direct comparison between short and long ITI conditions shows that the stimulation had a stronger impact on choices following long ITIs than choices following short ITIs in SERT-Cre mice, while there was no difference in WT mice (**Figure S11**).

In addition, in the absence of photo-stimulation, the correlation was smaller for the SERT-Cre mice than the WT mice (**Figure 2b**). This could indicate a chronic effect of stimulation [11], or a baseline effect of the genetic constructs, in addition to the trial-by-trial effect.

### 2.4 Optogenetic stimulation of DRN 5-HT neurons increased learning rate: Model-based analysis

Our analysis so far suggests that the choices following short ITIs follow a relatively simple win-stay lose-shift rule, while choices following long ITIs reflect a more gradual learning about reward and choice histories over multiple trials. Furthermore, we showed that optogenetic stimulation of 5-HT neurons influenced the impact of reward history on choices following long, but not short, ITIs.

In order to test these findings in a more integrated way, we built a combined characterization of choice. Figure 3a depicts a model in which there is an ITI threshold (now treated as a free parameter rather than being set to 7s) arbitrating whether the previously validated two-kernel model [39, 22] (i.e. short-term learning based win-stay lose-switch model), or a longer-term reinforcement learning (RL) model [56] would determine choice. The RL model allowed for two different learning rates associated with the prediction error on a given trial (**Figure 3b**): *α*_Stim_ (for stimulated trials) and *α*_No-Stim_ (for non-stimulated ones). We found that this model fits the data more proficiently than a number of variants (see the Methods section for details) embodying a range of dif-ferent potential effects of optogenetic stimulation: including acting as a direct reward itself; as a multiplicative boost to any real reward; or causing a change in the learning and/or forgetting rates (**Figure S8**).

**Figure 3:**
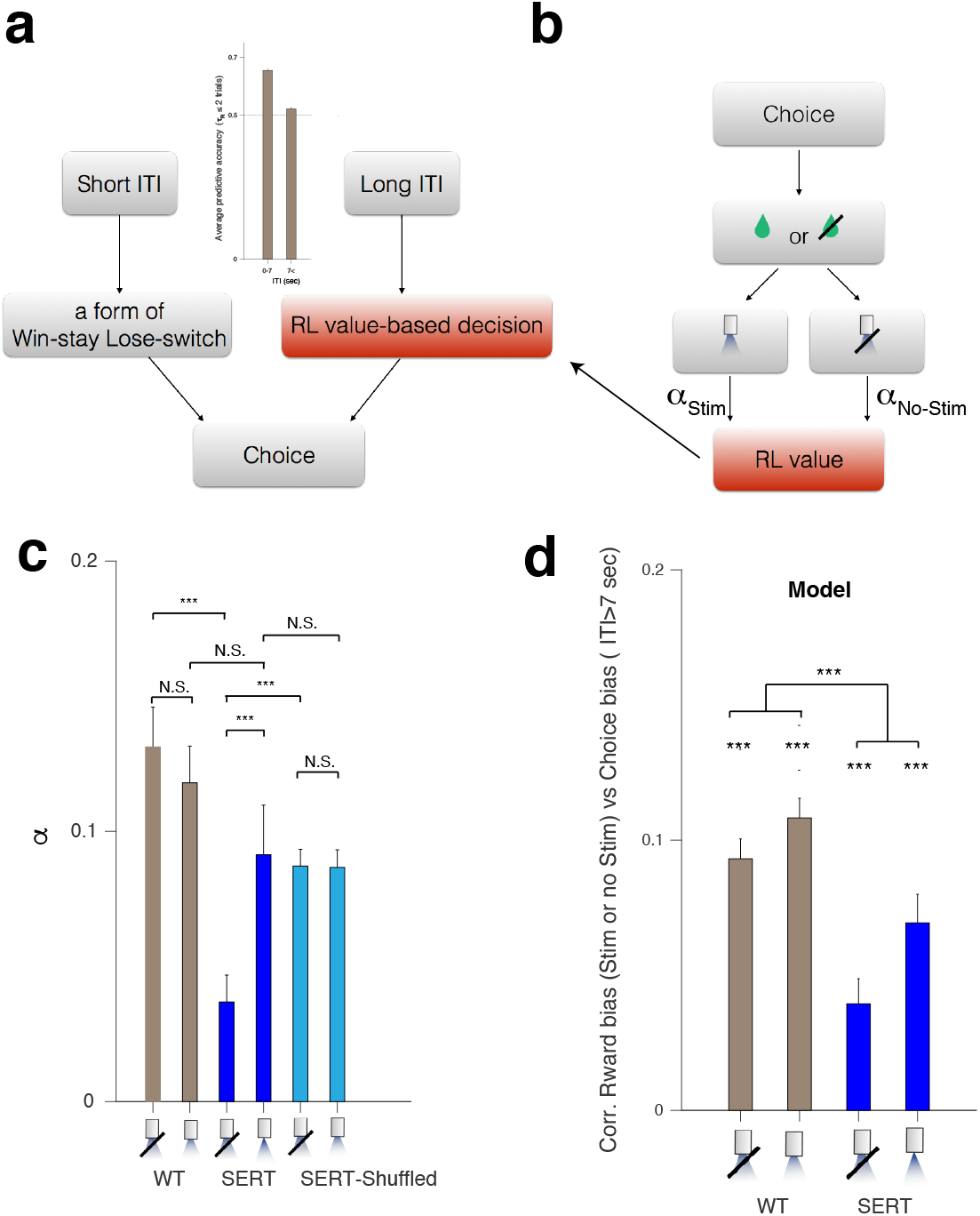
Photostimulation increased the learning rate of SERT-Cre mice. (**a**) Schematics of the computational model. There are two separate decision making systems: a fast system generating a form of “win-stay, lose-switch”, and a slow system following reinforcement learning (RL). After short ITIs *(T*_*ITI*_ < *T*_Threshold_), choice is generated by the fast system following win-stay, lose-switch. After long ITIs *(T*_*ITI*_ > *T*_Threshold_), choice is generated by the slow RL system. The ITI threshold *T*_Threshold_ is a free parameter that is fitted to data. (**b**) The RL system is assumed to learn the value of choice on all trials, including those with short ITIs for whose choices it was not responsible. The learning rate of the RL system is allowed to be modulated by photo-stimulation. When photo-stimulation is (respectively, is not) delivered, choice value is updated at the rate of *α*_Stim_ (*α*_no-Stim_). (**C**) Photostimulation increased the learning rate of SERT-Cre mice. The estimated learning rates for the WT (left), SERT-Cre (center), SERT-Cre mice (right) with shuffled stimulations are shown. The difference between *α*_*Stim*_ and *α*_no-Stim_ in WT mice, between *α*_Stim_ in WT mice and *α*_Stim_ in SERT-Cre mice, between *α*_Stim_ *α*_no-Stim_ in SERT-Cre mice with shuffled stimulation conditions, and between *α*_Stim_ in SERT-Cre mice and *α*_Stim_ in SERT-Cre mice with shuffled stimulation conditions were not significant. The difference between *α*_Stim_ and *α*_no-Stim_ in SERT-Cre mice, and between *α*_no-Stim_ in WT and *α*_no-Stim_ in SERT-Cre mice were significant (permutation test, *p <* 0.001). The difference between *α*_no-Stim_ in SERT-Cre mice and *α*_no-Stim_ in SERT-Cre mice with shuffled stimulations was also significant (permutation test, *p* < 0.01). (**d**) Generative test of the model. The analysis of Figure 2d was applied to data generated by the model. The correlations were all significantly differently from zero, while the difference between photo-stimulation and no photo-stimulation conditions between WT and SERT-Cre mice was also significant.

This model also fits choices better than a model that learns and forgets outcome history according to wall-clock time (measured in seconds) rather than according to the number of trials. To do this we simply adapted the previously validated two-kernel model that integrates choice and reward history over trials [39, 22] such that the influence of historical events is determined by how many seconds ago they happened, using the factual timing of the experiments. Model comparison using WT mice favored the account of **Figure 3a** (Δ iBIC = 218). Introducing two time constants to the reward integration kernel did not change this conclusion.

In the best fitting model (**Figure 3a**), we found that optogenetic stimulation increased the learning rate in SERT-Cre mice, but not in WT mice (**Figure3c**). Consistent with the previous analyses, we also found that the time constants for the choice kernel and the reward kernel for choices following short ITIs were very short for both WT and SERT-Cre mice (**Figure S5**), and that the ITI thresholds were not significantly different between WT and SERT-Cre mice (**Figure S6**). In addition, we replicated the same results using a model with a fixed (= 7 sec) ITI threshold (**Figure S7**).

As a control analysis, we fitted the model to SERT-Cre data with randomly reassigned stimulation trials. Shuffling the trials abolished the effect of photo-stimulation on the learning rate (**Figure 3c**).

Although the learning rate on stimulation trials in SERT-Cre mice was significantly greater than that on non-stimulated trials, it was not significantly different from the learning rate in WT mice (**Figure 3c**).), as already hinted by the model-agnostic analysis (Figure2b).

As hinted at by the model-agnostic analysis in **Figure2b**, the learning rate on no-stimulation trials was significantly smaller than that on stimulation trials in SERT-Cre mice.

Finally, we performed a generative test of the model to assess its ability to capture key aspects of the data. To do this, we simulated our model 100 times using each collection of parameters fit to each session of each subject, and analyzed generated data using the model agnostic procedures adopted for the original data (shown in **Figure2b**). As for the real data, the simulated data also showed a significant correlation between reward history and the choice-after-long-ITIs, and a significant difference between photo-stimulation and no photo-stimulation conditions between WT and SERT-Cre mice (**Figure 3d**).

Our analysis has so far focused on the impact of reward history over a relatively short timescale (< 50 trials) compared to the length of a whole experimental session (> 100 trials). Since animals can also learn reward histories over much longer timescales [10, 31], and 5-HT neurons have shown to encode reward rates over multiple timescales [8], it is possible that the optogenetic stimulation of DRN neurons might have had effects over hundreds of trials. To examine this, we conducted a simple correlation analysis by dividing each session into five quintiles (containing equal numbers of trials), as in **Fig. S3** and asked how the choices following long ITIs in the last quintile (the only one with substantial numbers of long ITI choices) were correlated with the reward history stretched over all numbers of preceding quintiles (e.g. only the fifth, the fourth and the fifth, etc.). For reward history, we used the probabilities determined by the experimenters rather than those observed by the subjects, to avoid any bias that is independent of the reward history (such as choice history).

Choices following long ITIs were indeed significantly influenced by long run reward history spanning over the entire experimental session (**Figure S9**). The data from the generative test also confirms this correlation (**Figure S9**), albeit to a lesser degree, perhaps because the model only involves a single time constant and may thus have an inflated learning (and thus forgetting) rate relative to these long gaps. Furthermore, although the data shows that these effects were stronger in SERT-Cre mice than in WT mice (2-way ANOVA; *p =* 0.0016, *F* = 11.98), we did not see this in our generative test results. Thus longer time constants (slower learning) that are present [10, 30, 31] may also be affected by genotype or actual optogenetic stimulation.

## 3 Discussion

There have been many suggestions for the roles that serotonin might play in decision-making and choice. These include ideas about influences over motor behavior [35], punishment [18, 19, 51], opponency with dopamine [21, 13, 6], satiation [57], discounting [20] patience [45, 22] and even aspects of reward [42, 8, 44]. Here, we report an additional effect: serotonin stimulation can increase the rate at which animals learn from choice outcomes in dynamic environments.

A standard learning rule in RL has two distinct components. The first is the reward prediction error (RPE), which quantifies the difference between the actual and predicted value of outcomes. The phasic activity [46] of midbrain dopamine neurons and the local concentration of dopamine [26, 37] in target regions follow this pattern. The second component is the learning rate, which determines how much change is actually engendered by the prediction error. From a normative perspective, learning rates are determined by the degree of uncertainty [16] – influenced by factors such as initial ignorance and the volatility of the environment, since we should only learn when there is something that we do not know. There is experimental evidence that this is indeed the case [24, 48, 4, 47]. While it has been suggested that the neuro-modulators norepinephrine (NE) and acetylcholine (ACh) may influence learning rates [62, 2], our findings suggest that 5-HT DRN neurons also play a critical role. The interaction between 5-HT and dopamine could potentially be implicated in this effect, as various serotonin receptor types can increase the release of dopamine [17], which, if operating at an appropriate timescale, could boost the effective learning rate.

It is notable that the effect of altered learning rates was only apparent on trials following long than short ITIs. The former choices also hewed to a different strategy than the latter. Short ITIs appeared to lead to decisions closer to win-stay, lose-shift, meaning that subjects weighed barely more than the outcome of the most recent trial in their decision. The shift between strategies might correspond to a difference between a policy based on working memory [9] for very recent events (a few seconds) vs. a plasticity-based mechanism like that assumed by standard incremental RL for incorporating events over longer periods. Note, though, that the boundaries between working memory and RL are becoming somewhat blurred [5]. It has been suggested that memory based methods contribute to model-based control, by contrast with incremental model-free RL [41, 14, 40, 5]; but this remains to be pinned down experimentally. Note that a similar effect has been also observed in perceptual decision making. In one example, longer-lasting prior experience was more influential when working memory was disturbed during the task [1].

The distribution of short and long ITI trials suggests that they might reflect the animal’s motivational state as being high and low, respectively. Long ITI choices were most frequent in the last quintile of each experimental session, where animals were likely to be sated. That they also occurred in the beginning of experimental sessions might suggest that the subjects were not fully engaged in the task at the start, perhaps hoping to get out from the experimental chamber. A more systematic analysis of behavior during long ITIs would be required to uncover the nature of those events.

The fact that only a subset of trials was apparently affected by the stimulation is arguably a cautionary tale for the interpretation of optogenetics experiments. What looked like a null effect [22] had to be elucidated through computational modeling. Equally, for the short ITI trials, what seemed like behavior controlled by conventional RL, might come from a different computational strategy (and potentially neural substrate) altogether [9]. This could prompt a reexamination of previous data (as shown by [5]). Further caution might be prompted by the observation that the learning rate in the SERT-Cre mice in the absence of stimulation was actually significantly lower than that of the WT mice in the absence of stimulation, rising to a similar magnitude as the WTs, with stimulation. This may be due to chronic effects of optogenetic stimulation of DRN neurons, as suggested in recent experiments [11], or due to baseline effects of the genetic constructs.

The learning rates that we found even for the slow system are a little too fast to capture fully the long term correlation that can be found in the data. This is apparent in our additional analysis showing the correlation between the reward bias in the 1st quintile of the trials and the choice bias in the 5th quintile of the same session (**Figure S12**), also the correlation between the reward bias in the 5th quintile of the trials and the choice bias in the 1st quintile in the succeeding experimental session (**Figure S13**). The former is surprising, since it spans a large number of trials; the latter because it usually spans more than a day. This could suggest that learning in fact took place over a wide range of timescales, and the time constant that we found by our model-fitting reflects a weighted average of those multiple time constants [30, 31]. It would be interesting to study how the duration of ITI, or the level of engagement in the task, can change the weight or relative contribution of those distinctive time constants. It is plausible that the two decision strategies that we considered here are just an approximation to a wider collection of strategies that operate over a wider range of distinctive timescales. It would then be interesting to ask why serotonin stimulation preferentially affected slower components. Further questions include whether serotonin’s effects would be better captured as an influence on the relative weighting of different timescales [31] rather than the changes in time constants themselves that we assumed in the model fitting.

Finally, one of the main reasons to be interested in serotonin is the prominent role that drugs affecting this neuromodulator play in treating psychiatric disorders. While our results add substantial complexity to this landscape, they also offer the prospect of richer and more finely targeted manipulations, given greater understanding.

## Acknowledgement

We thank the Gatsby Charitable Foundation, the Joint Initiative on Computational Psychiatry and Ageing Research between Max Planck Society and UCL, the Japan Society for the Promotion of Science, the European Research Council (250334 and 671251), Fundação para a Ciência e a Tecnologia (PD/BD/52446/2013 and f SFRH/BPD/46314/2008) and the Champalimaud Foundation for generous support.

## Methods

### M.1 Inter-trial-interval (ITI)

We defined the inter-trial-interval (ITI) as the time from when the mouse left one of the side ports until it entered the center port to initiate the next trial. Occasionally, animals re-visited the side port long after their first visit on the same trial (less than 5 per ent of all trials). These redundant pokes were ignored.

### M.2 Computational models for decision making

#### M.2.1 Reward and choice kernel model [39, 22]

Previous studies have shown that animal’s choice behavior in a dynamic foraging task without the change-over-delay constraint [28] can be well-described by a linear two-kernel model (e.g. [39, 22]). In this model, the probability 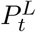 of choosing Left on trial *t* is determined by a linear combination of values computed from reward and choice history, given by

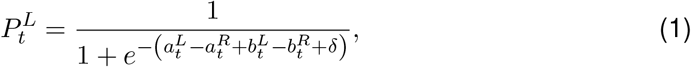

where 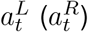 is the value computed from a reward kernel for Left (Right), 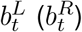 is the value computed from a choice kernel for Left (Right), and *δ* is the bias. Assuming simple exponential kernels [39, 54, 10], the reward values are updated on every trial as:

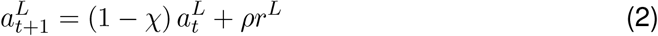

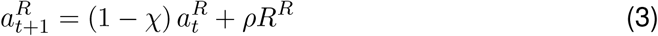

where 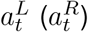 is the reward value for Left (Right) choice on trial *t, χ* is the temporal forgetting rate of the kernel, ρ is the initial height of the kernel, and *r*^*L*^ = 1 (*r*^*R*^ = 1) if a reward is obtained from Left (Right) on trial t, or 0 otherwise. Since these equations can also be written as:

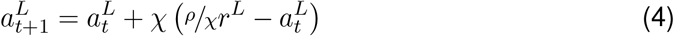

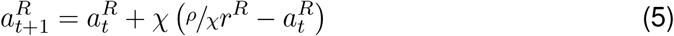

this kernel is equivalent to a forgetful Q-learning rule [60, 9] with a learning rate *χ* and reward sensitivity *ρ/χ*

The value for choice is also updated as

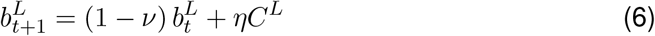

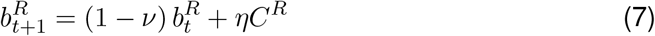

where 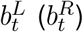 is the choice value for Left (Right) choice on trial *t*, *ν* is the temporal forgetting rate of the kernel, *η* is the initial height of the kernel, and *C*^*L*^ = 1 (*C*^*R*^ = 1) if Left (Right) is chosen on trial t while 0 otherwise. We note that the initial height of the choice kernel, *η*, is normally negative [39, 22], meaning that the choice kernel normally captures a tendency towards alternation. Such tendencies are common in tasks with reward schedules like those in the current task if a penalty for alternation is not imposed (change over delay) [28].

We assumed that the update takes place on every trial, even those associated with long ITIs.

#### M.2.2 Main model

We constructed a model that describes choices on all trials. Since we found that the characteristics of decision strategies changed according to the ITIs, we simply assumed a two-agent model, where agent 1 (fast system) makes decisions on the trials following short ITIs (ITI ≤ *T*_Threshold_), while agent 2 (slow system) makes decisions on the trials following long ITIs (ITI > *T*_Threshold_). We allowed the threshold *T*_Threshold_ to be a free parameter that is determined by data. We also tested the fixed value *T*_Threshold_ = 7 seconds based on our preliminary analyses and found results consistent with the variable ITI-threshold model (**Fig. S7**).

The fast system generates decisions based on the two-kernel model described in M.2.1. The slow system performs simple Q-learning. Specifically, the probability 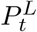 of choosing Left on trial **t** after a long ITI > *T*_Threshold_ is given by

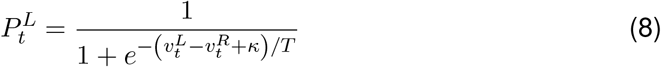

where 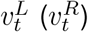 is the value for Left (Right), *κ* is the bias term, and **T** is the decision noise.

The agent updates values for chosen action according to the Rescorla-Wagner rule, but at different learning rates for photo-stimulation (*α*_Stim_) and no-stimulation (α_No-Stim_) trials. For example, if Left was chosen and photo-stimulation was applied, the value of Left choice is updated as

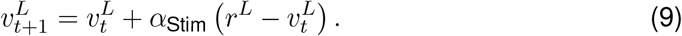

If no stimulation was applied, on the other hand,

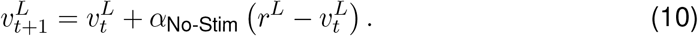

By comparison with equation 5, we can see this as a non-forgetful Q-learner, but with a slightly more convenient parameterization for the reward sensitivity. For a model comparison purpose, we also fitted a forgetful Q-learner model with optogenetically modulated learning rates, in which the updates given by Equations 9 and 10 take place for the values of both choices every trial.

Both systems updates values every trial regardless of the preceding ITIs, but the decision was made by one of them depending on the most recent ITI, where the threshold *T*_Threshold_ was also a free parameter. **Figure 3** shows the results of this full model.

#### M.2.3 Other models

In order to explore other possibilities for optogenetic stimulation effects, we constructed three other models.

##### Asymmetric learning rate model

We allowed the model to have different learning rates for reward and no-reward trials when photo-stimulation was applied. Specifically, we modified Equation 9 of the main model as

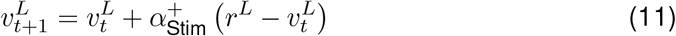

if *r*^*L*^ = 1, and

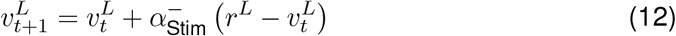

if *r*^*L*^ = 0. The same is applied for the Right choice.

##### Multiplicative value model

Here we assumed that photo-stimulation changed the sensitivity of reward. Specifically, we modified the learning rules of slow system as

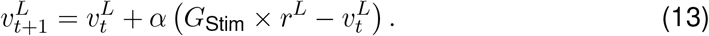

if photo-stimulation is applied, otherwise

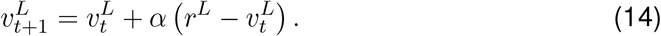

##### Additive value model

Here we assumed that photo-stimulation carried a independent rewarding value. Specifically, we modified the learning rules of slow system as

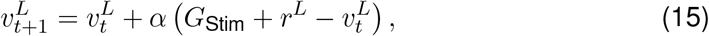

if photo-stimulation is applied, otherwise

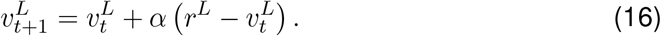

The same is applied for the Right choice.

### M.3 Model fitting

In order to determine the distribution of model parameters **h**, we conducted a hierarchical Bayesian, random effects analysis [29, 34, 33] for each subject. In this, the (suitably transformed) parameters **h**_*i*_ of experimental session *i* are treated as a random sample from a Gaussian distribution with means and variance 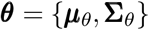.

The prior distribution ***θ*** can be set as the maximum likelihood estimate:

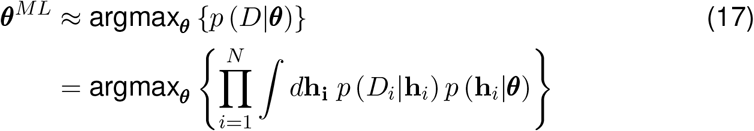

We optimized ***θ*** using an approximate Expectation-Maximization procedure. For the E-step of the k-th iteration, a Laplace approximation gives us

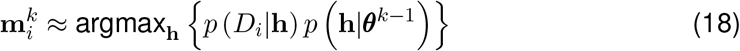

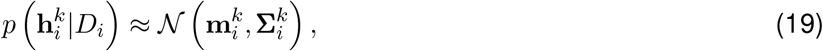

where 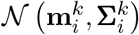 is the Normal distribution with the mean 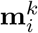 and the covariance 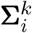 that is obtained from the inverse Hessian around 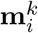. For the M step:

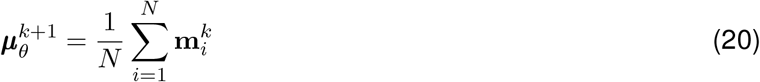

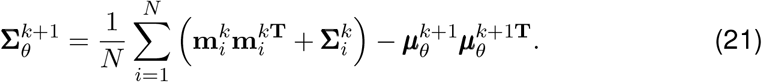

For simplicity, we assumed that the covariance 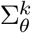 had zero off-diagonal terms, assuming that the effects were independent.

## Model comparison

We compared models according to their integrated Bayes Information Criterion (iBIC) scores [29, 34, 33]. We analysed model log likelihood log *p(D|M*):

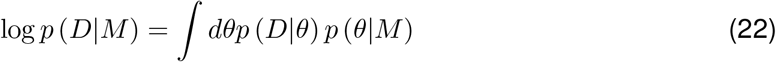

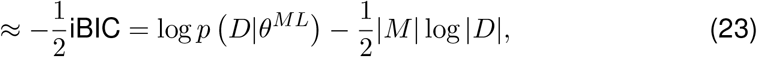

where iBIC is the *integrated* Baysian Information Criterion, |*M*| is the number of fitted prior parameters and |*D*| is the number of data points (total number of choice made by all subjects). Here, log *p*(*D*|*θ*^*ML*^) can be computed by integrating out individual parameters:

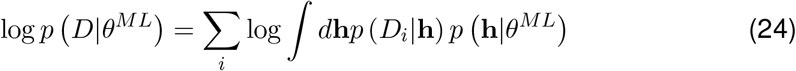

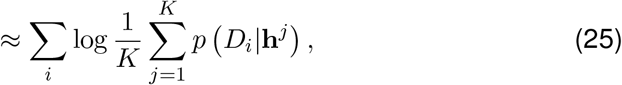

where we approximated the integral as the average over *K* samples **h**^*j*^’s generated from the prior *p* (**h**|*θ*^*ML*^).

## Model’s average predictive accuracy

We defined the model’s average predictive accuracy as the arithmetic mean of the likelihood per trial, using each session’s MAP parameter estimate. That is,

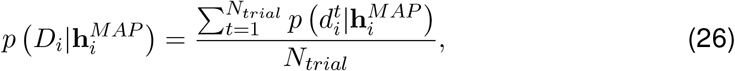

where *N*_*trial*_ is the number of the trial, 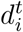 is the datapoint on trial *t* in session *i*.

In our generative simulations, we used the same reward/photo-stimulation schedule as the actual data.

## S1 Supporting figures

**Figure S1:**
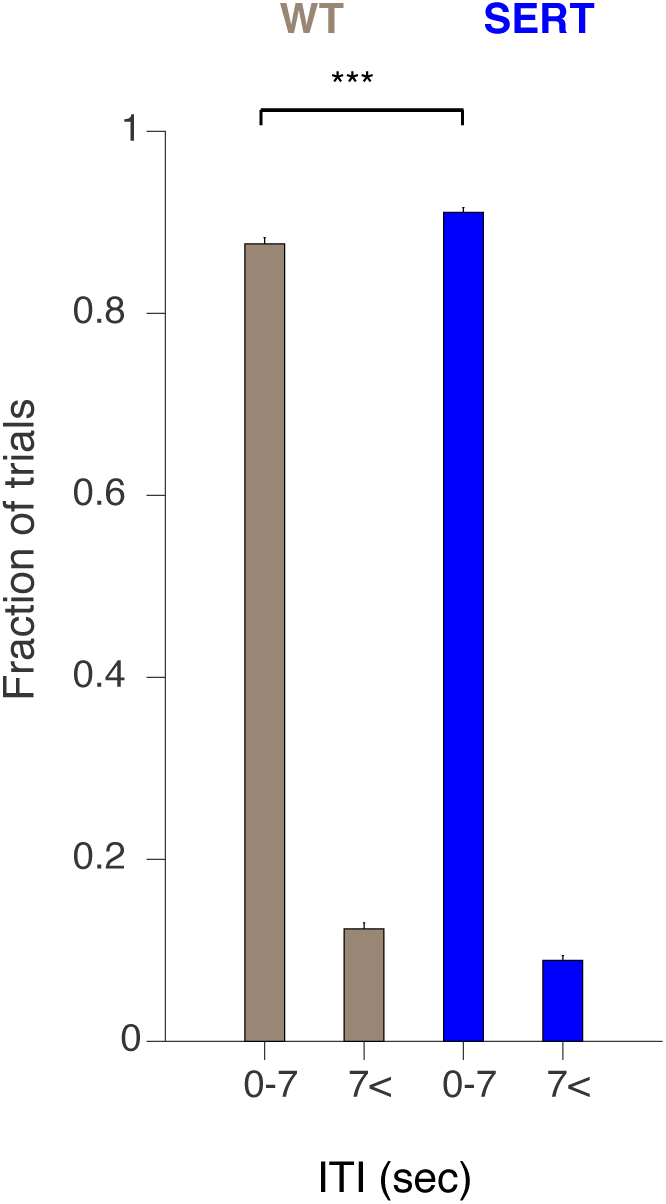
The distribution of ITIs. The proportions of short (≤ 7 sec) ITI trials and long (> 7 sec) ITI trials were significantly different for both WT (left) and SERT-Cre (right) mice. The difference between WT and SERT-Cre mice was also significant, though the optogenetic stimulation itself did not change the subsequent ITIs (see **Figure S2**). The error bars indicate the mean ± SEM of sessions.

**Figure S2:**
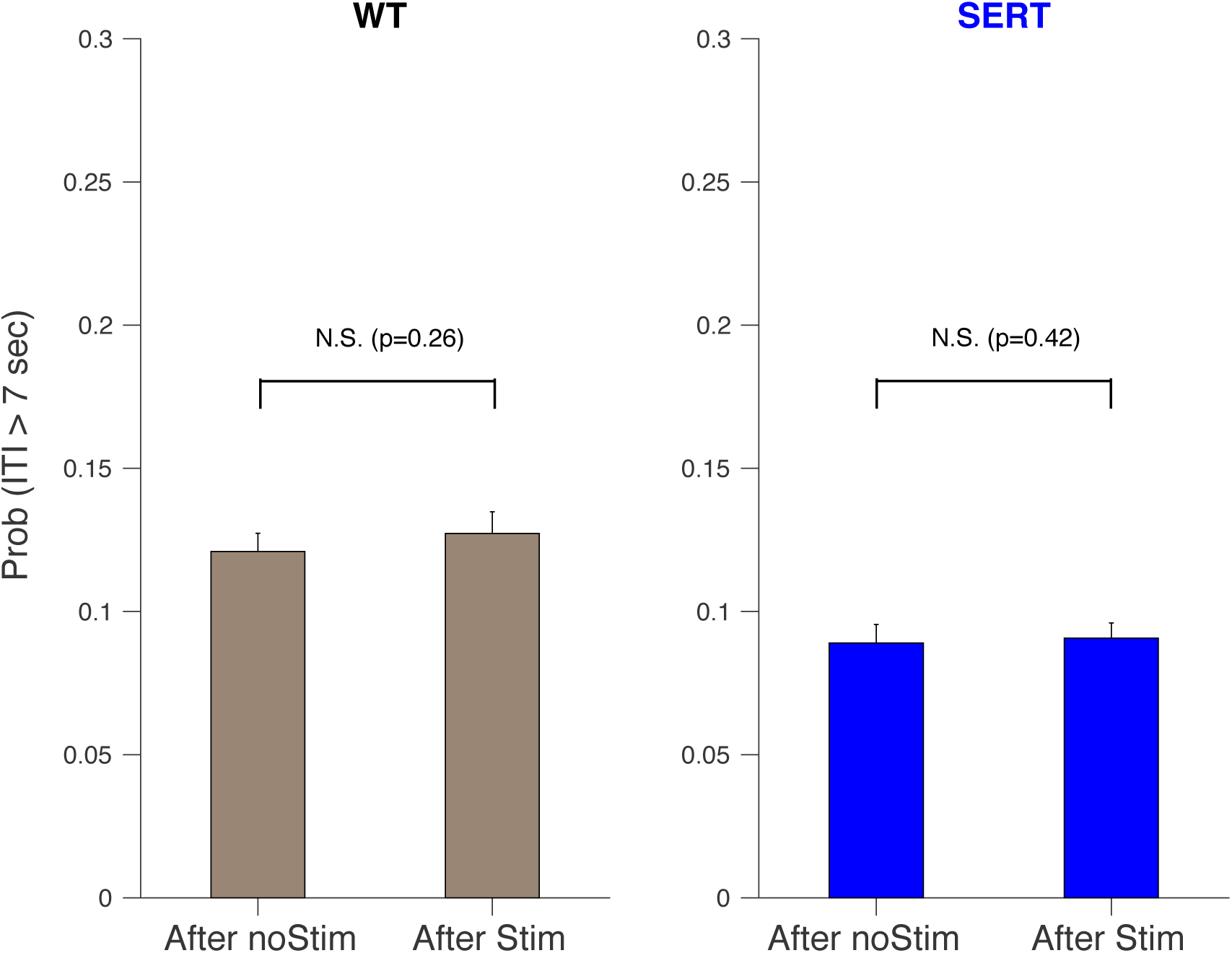
Probability that the ITI is longer than 7 sec, following a photo-, or no photo-, stimulation. Stimulation does not significantly increase the chance of creating a long ITI event.

**Figure S3:**
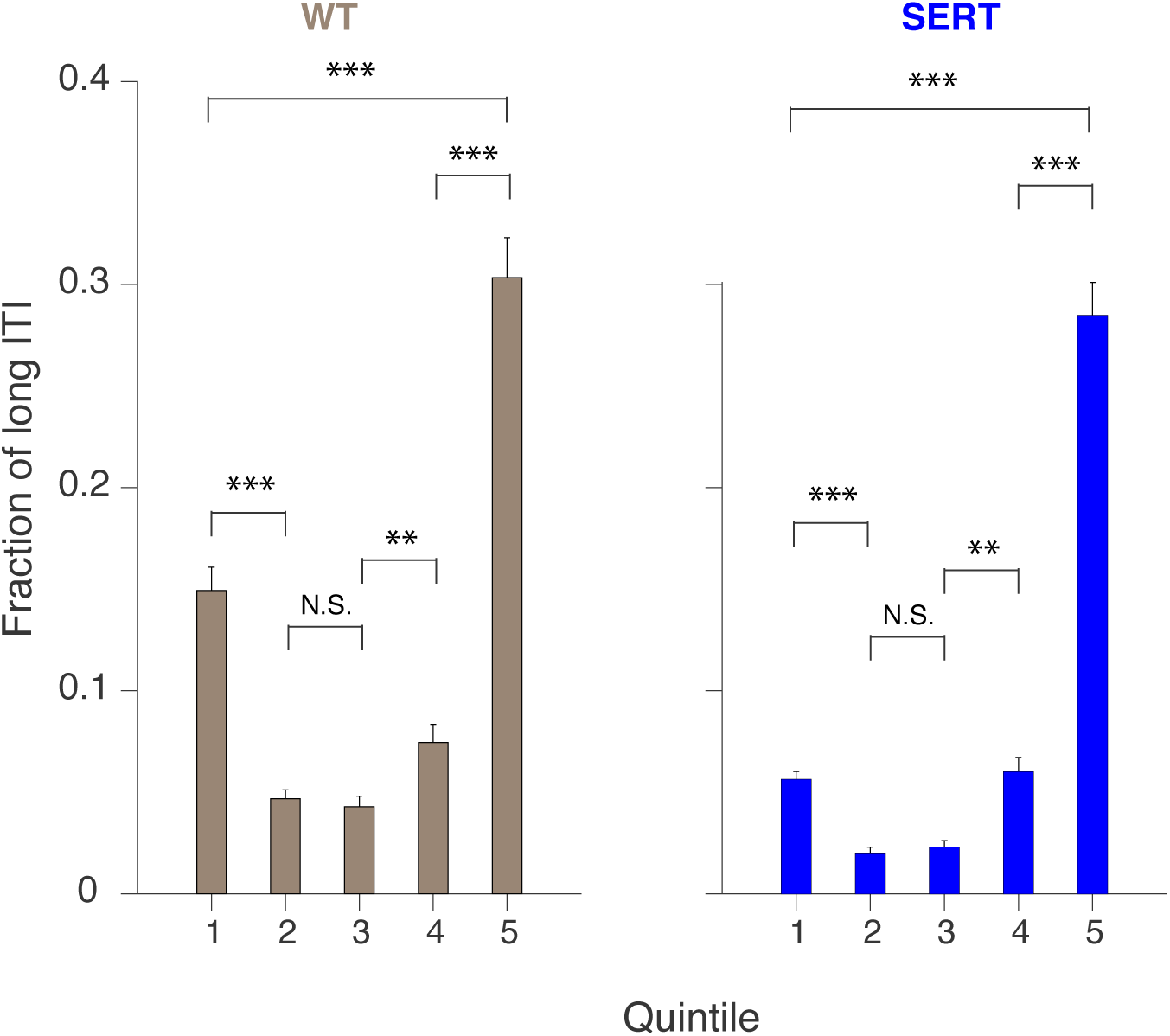
The fractions of long ITI trials in quintiles (containing equal numbers) of trials within sessions for wild-type (left; grey) and SERT (right; blue) mice. The error bars indicate the mean ± SEM. The difference between the first and the second quintile *(p* < 0.001, permutation test), between the third and the fourth quintile (*p* < 0.01, permutation test), between the fourth and the fifth quintile (*p* < 0.001, permutation test), between the first and the fifth quintile (*p* < 0.001, permutation test) are significant within WT and SERT-Cre mice, respectively.

**Figure S4:**
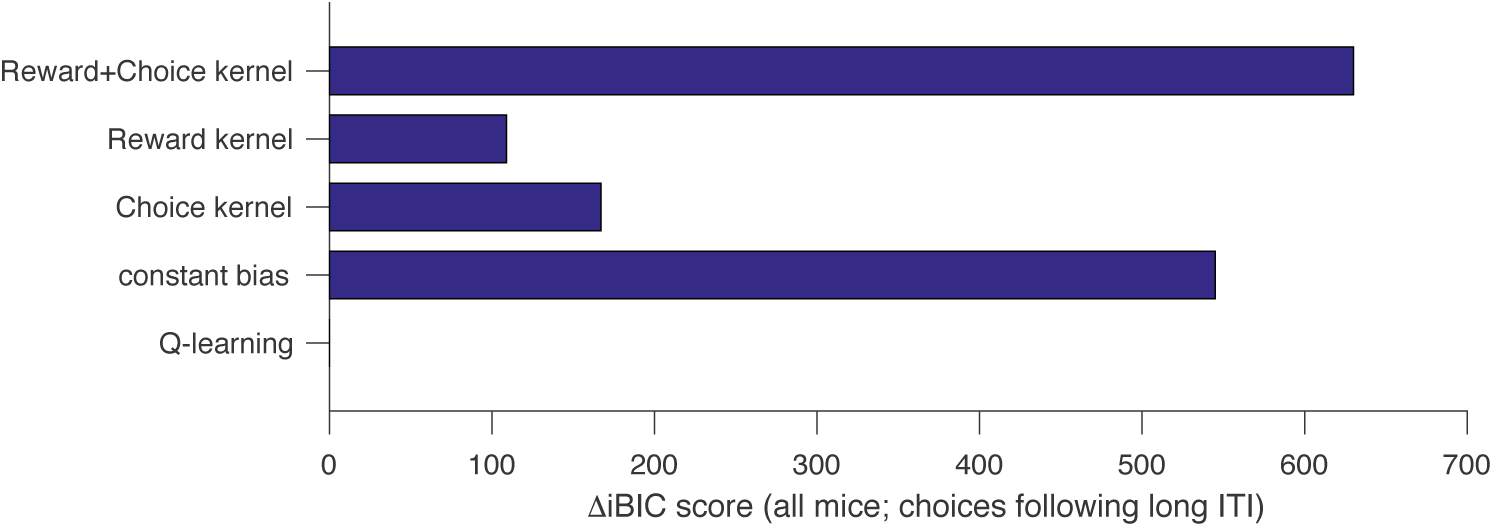
Model comparison for choices following long ITIs, based on integrated Bayesian Information Criterion (iBIC). Q-learning model outperforms the other models. The previously validated model, Reward + Choice kernel model (top), performs poorly for choices following long ITIs.

**Figure S5:**
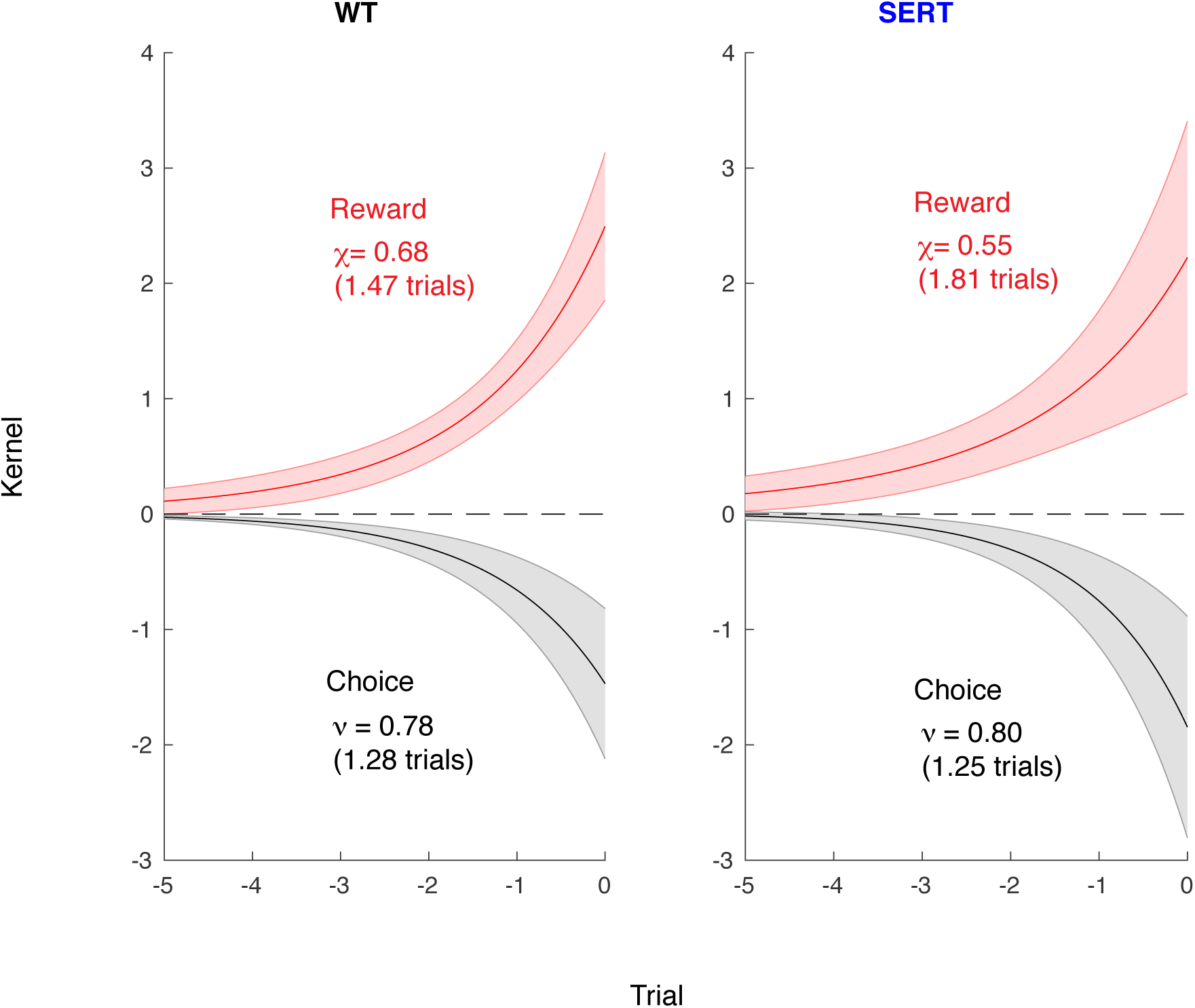
Estimated choice kernel and reward kernel for the fast system in the full model. The mean ± standard deviation of estimated kernels of all traces are shown for WT (left) and SERT-Cre (right) mice. The time constants are the mean of the estimates, not the re-fit of the mean trace.

**Figure S6:**
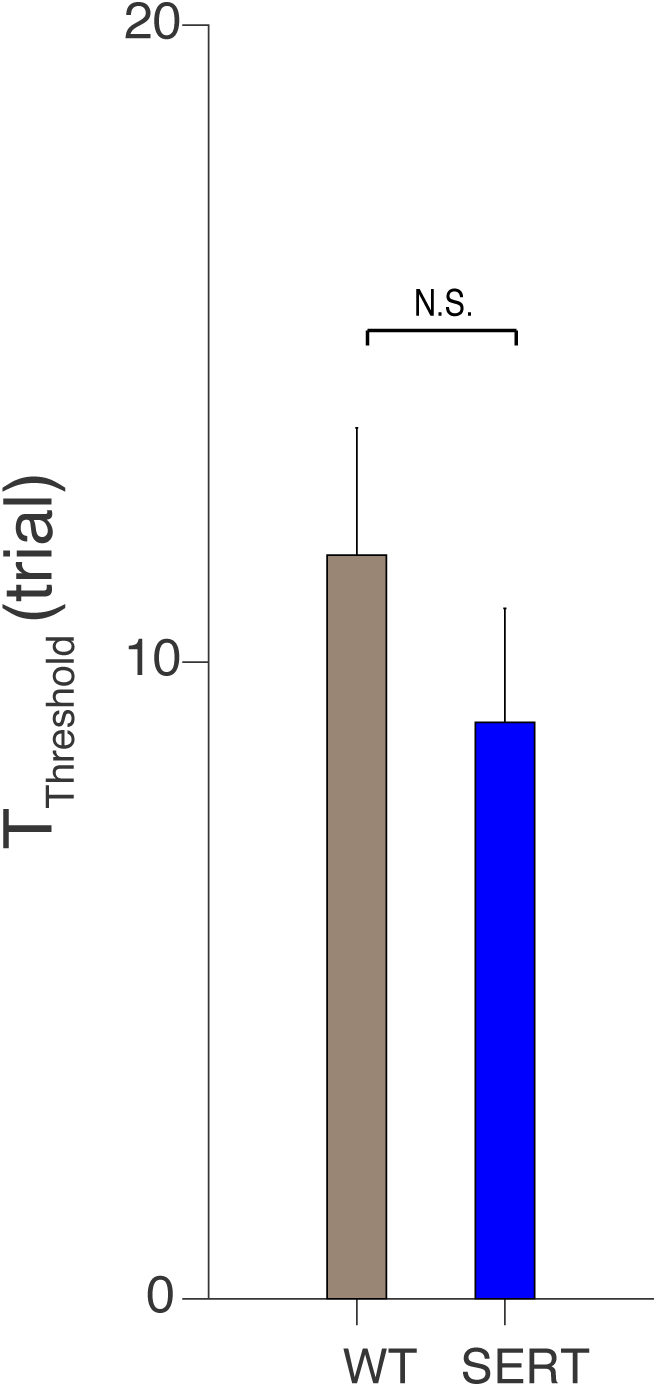
Estimated threshold for the full model. The mean ± SEM of estimated kernels are shown for WT (left) and SERT-Cre (right) mice.

**Figure S7:**
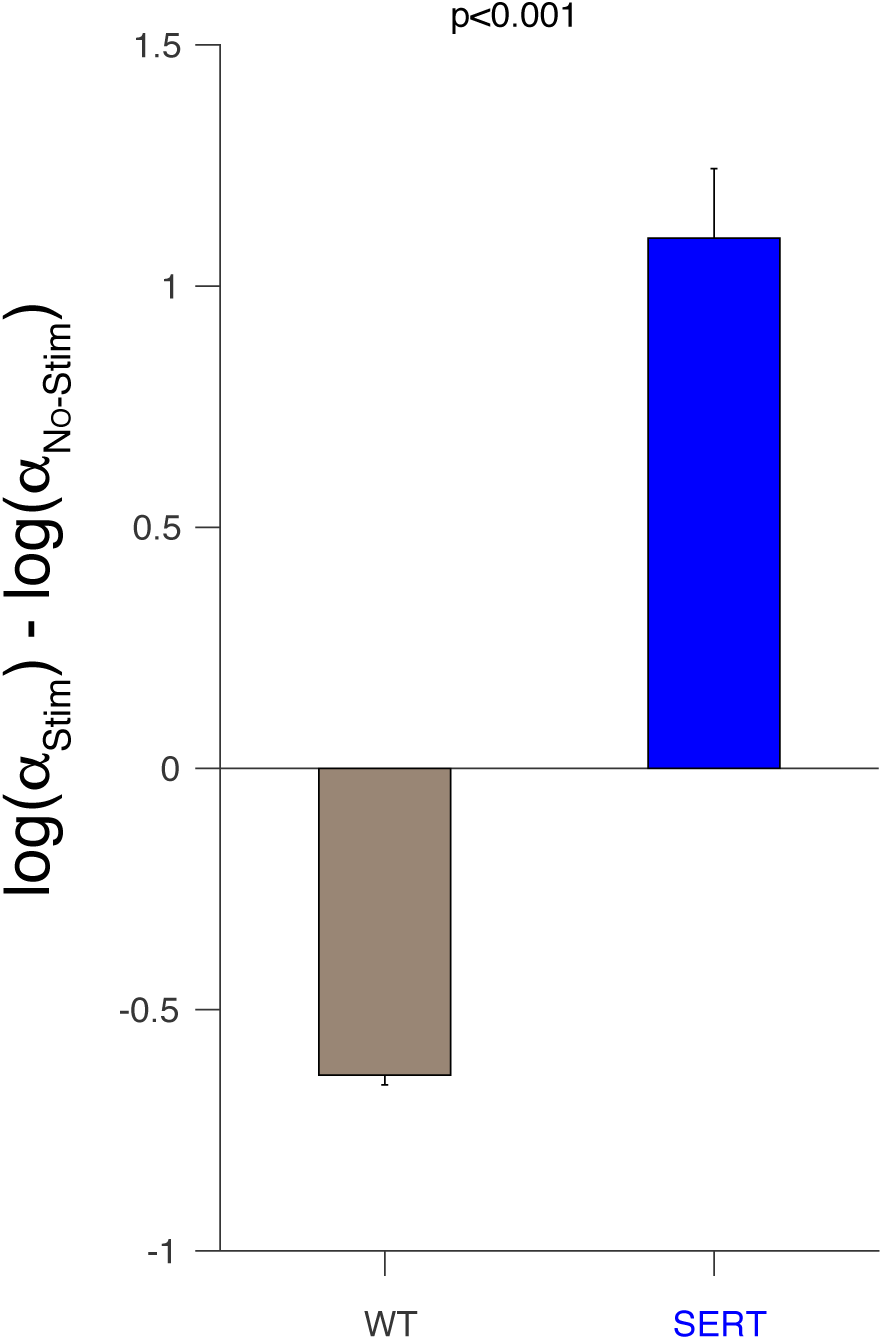
[Fixed-threshold model]. Photo stimulation increased the learning rate in SERT-Cre mice. The difference between the WT mice and SERT-Cre mice was significant (permutation test, *p <* 0.001). The simple Q-learning model was assumed to learn values on all trials but was responsible for decisions on trials following long ITIs (> 7 sec).

**Figure S8:**
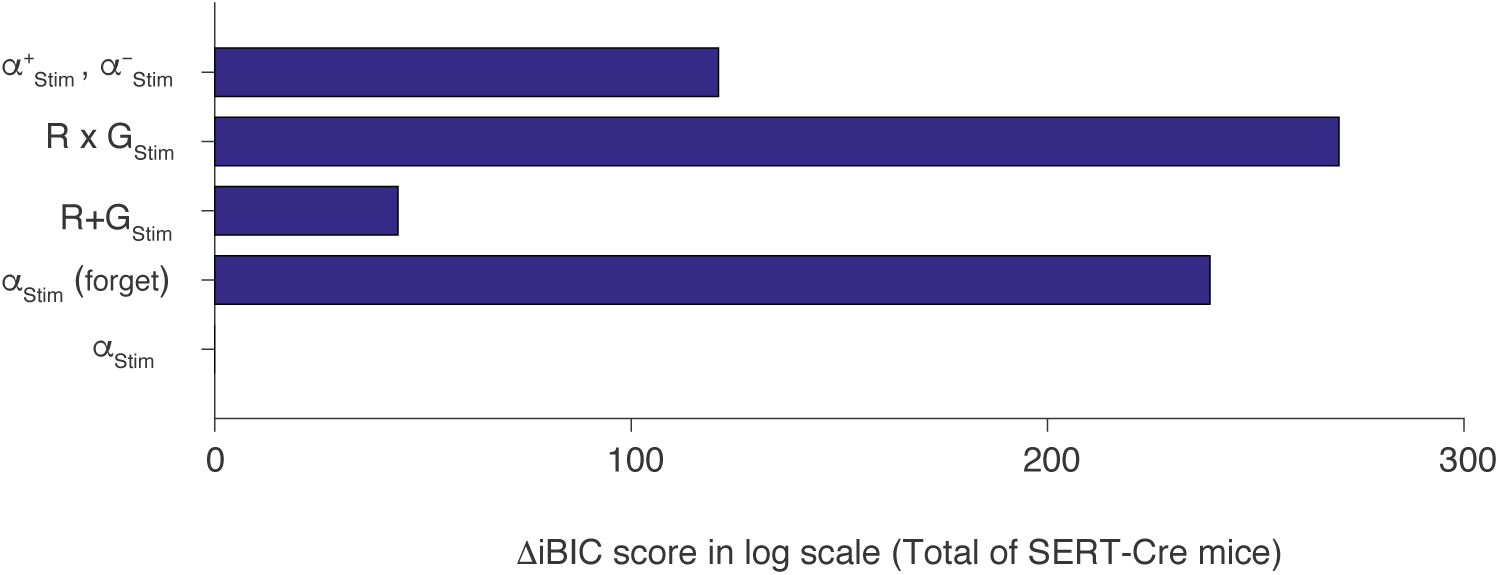
Model comparison for different optogenetic stimulation effects in SERT-Cre mice, based on integrated Bayesian Information Criterion (iBIC). Our model with a modulated learning rate (bottom) outperformed models with asymmetrically modulated learning rate (top), multiplicatively modulated reward value (2nd row), and additively modulated reward value (3rd row).

**Figure S9:**
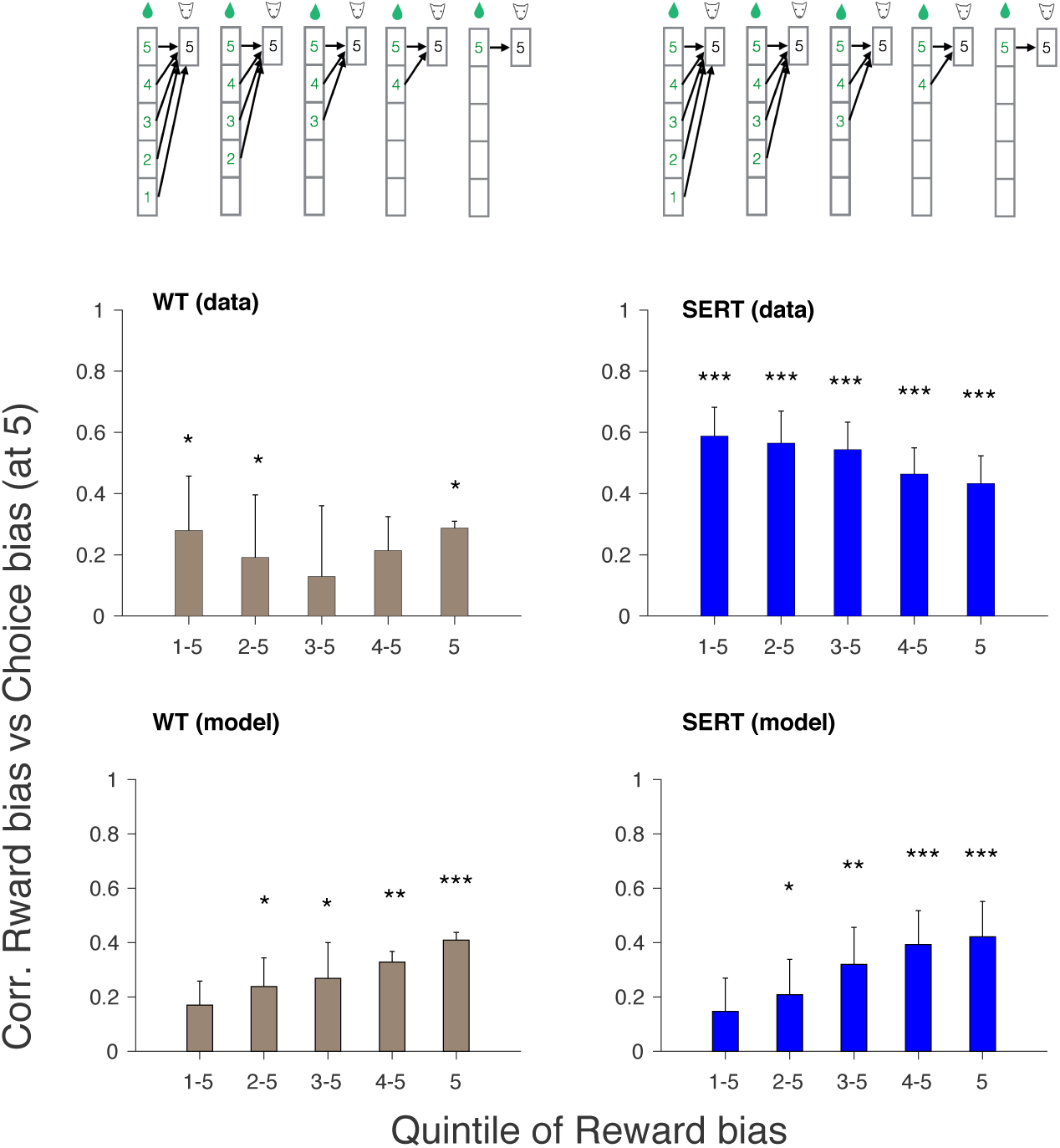
Choices following long ITIs were predicted by reward history over many trials. The correlation between the choices following long ITIs in the last part of each session (5th quintile) and the reward bias estimated in different quintiles. On the x-axis, ‘1-5’ indicates the overall reward bias computed in the total of the 1st, 2nd, 3rd, 4th and 5th quintiles, while ‘5’ means the bias from the 5-th quintile only. The top left (top right) shows the results of WT (SERT-Cre) mice, while the bottom left (bottom right) shows the results of model’s generated data for WT (SERT-Cre) mice. The stars indicate how significantly the correlation is different from zero, tested by a permutation test. The test statistic was constructed by the mean of the correlation coefficients of four animals at each quintile condition, where the correlation coefficient was computed by randomly permuted data in each condition in each animal. One star indicates *p <* 0.05; two stars indicates *p <* 0.01, while three stars indicates *p* < 0.001. The error bars indicate the mean ± SEM of data.

**Figure S10:**
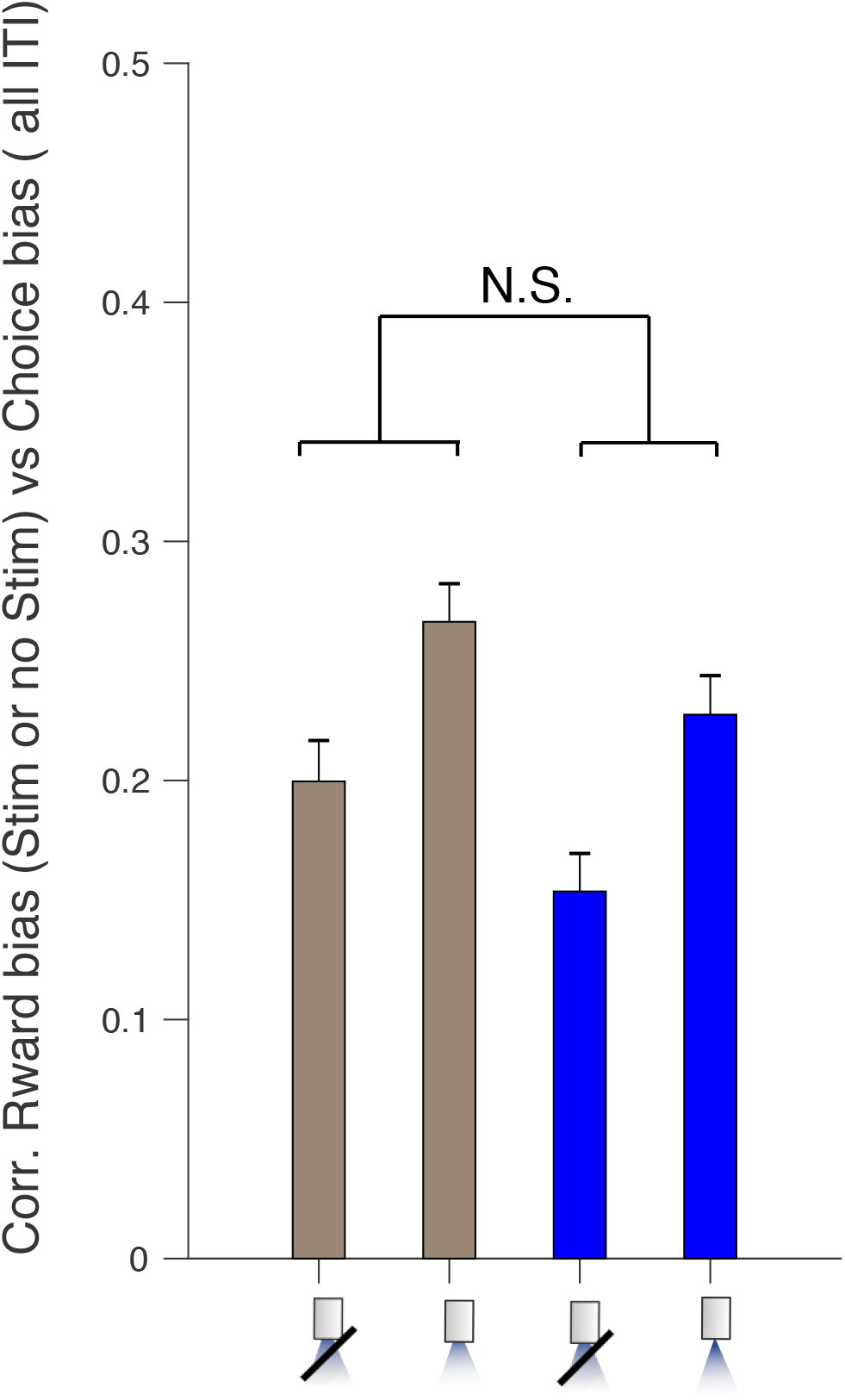
The impact of reward history on choices following short ITIs did not show effects of optogenetic stimulation. The x-axis indicates if the reward bias was computed over trials with or without photo-stimulations. Due to the experimental bias of stimulation and reward probability, the correlation appears to be larger when stimulation is on for both groups; however, the difference between WT and SERT-Cre was not significant. The error bars indicate the mean ± SEM of sessions.

**Figure S11:**
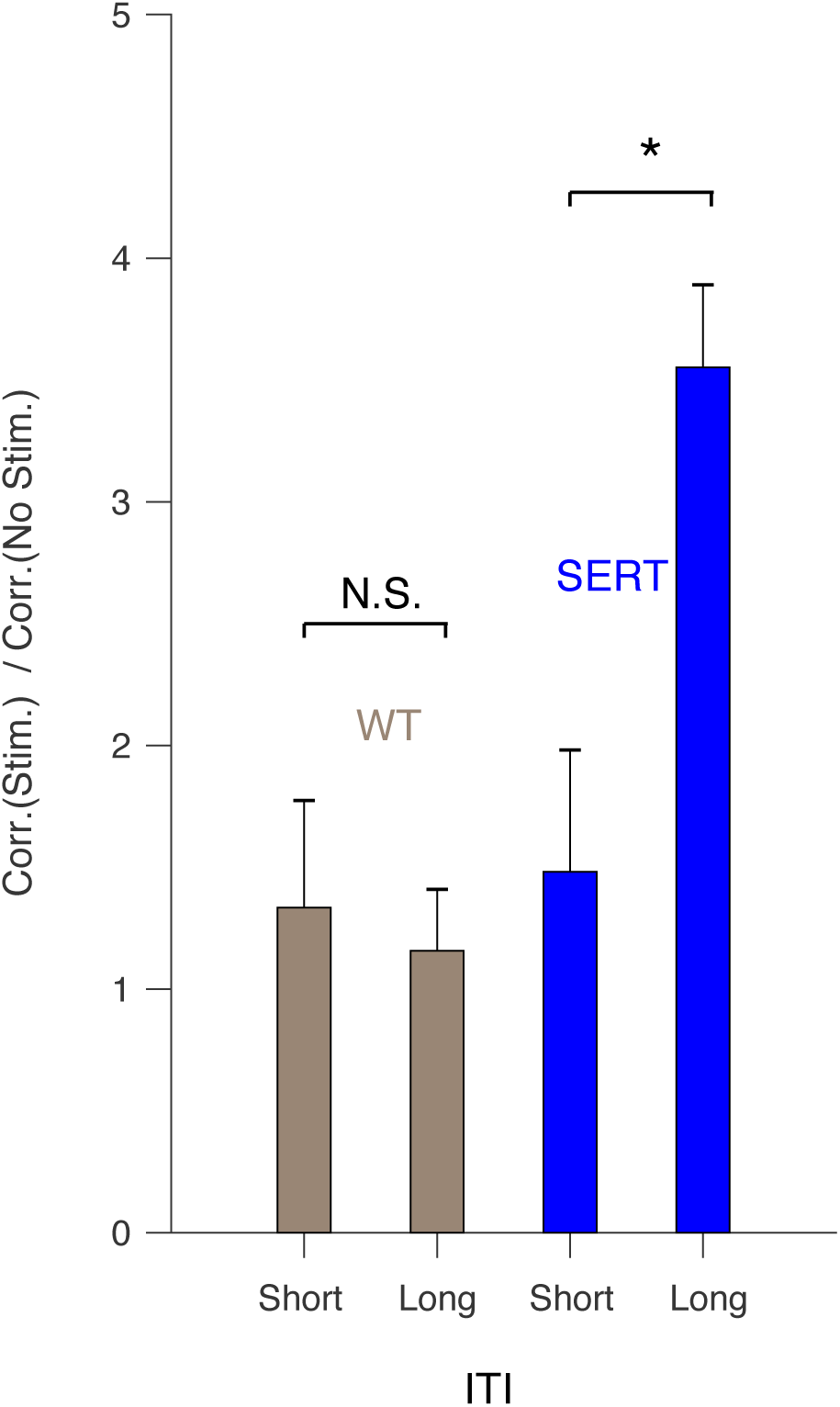
The impact of reward history on choice was more strongly seen in choices following long ITIs than ones following short ITIs in SERT-Cre mice, while it was not the case in WT mice. The x-axis indicates if the correlation was computed for choices following short or long ITIs. The y axis indicates the ratio of the correlation between reward bias and choice bias computed over trials with photo-stimulations to the correlation computed over trials without photo-stimulations. The difference between the short and the long ITI conditions in SERT-Cre mice was significant (permutation test; *p <* 0.05), while the difference in WT mice was not significant. The error bars indicate the mean ± SEM of data.

**Figure S12:**
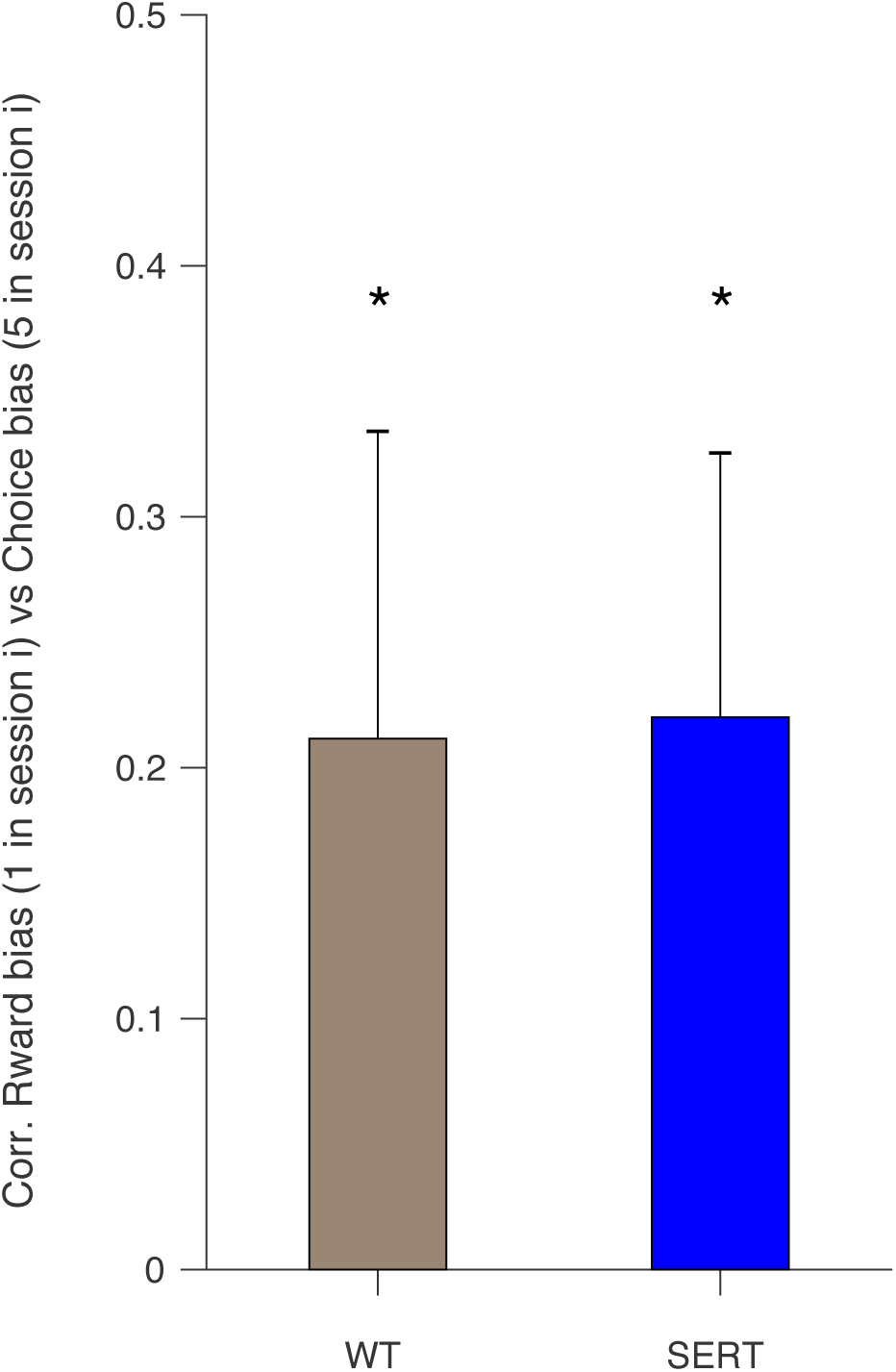
Choices following long ITIs in the fifth quintile were correlated with reward bias in the first quintile in the same session. The star indicate how significantly the correlation is different from zero, tested by a permutation test. The test statistic was constructed by the mean of the correlation coefficients of four animals, where the correlation coefficient was computed by randomly permuted data in each animal. One star indicates *p* < 0.05. The error bars indicate the mean ± SEM of data.

**Figure S13:**
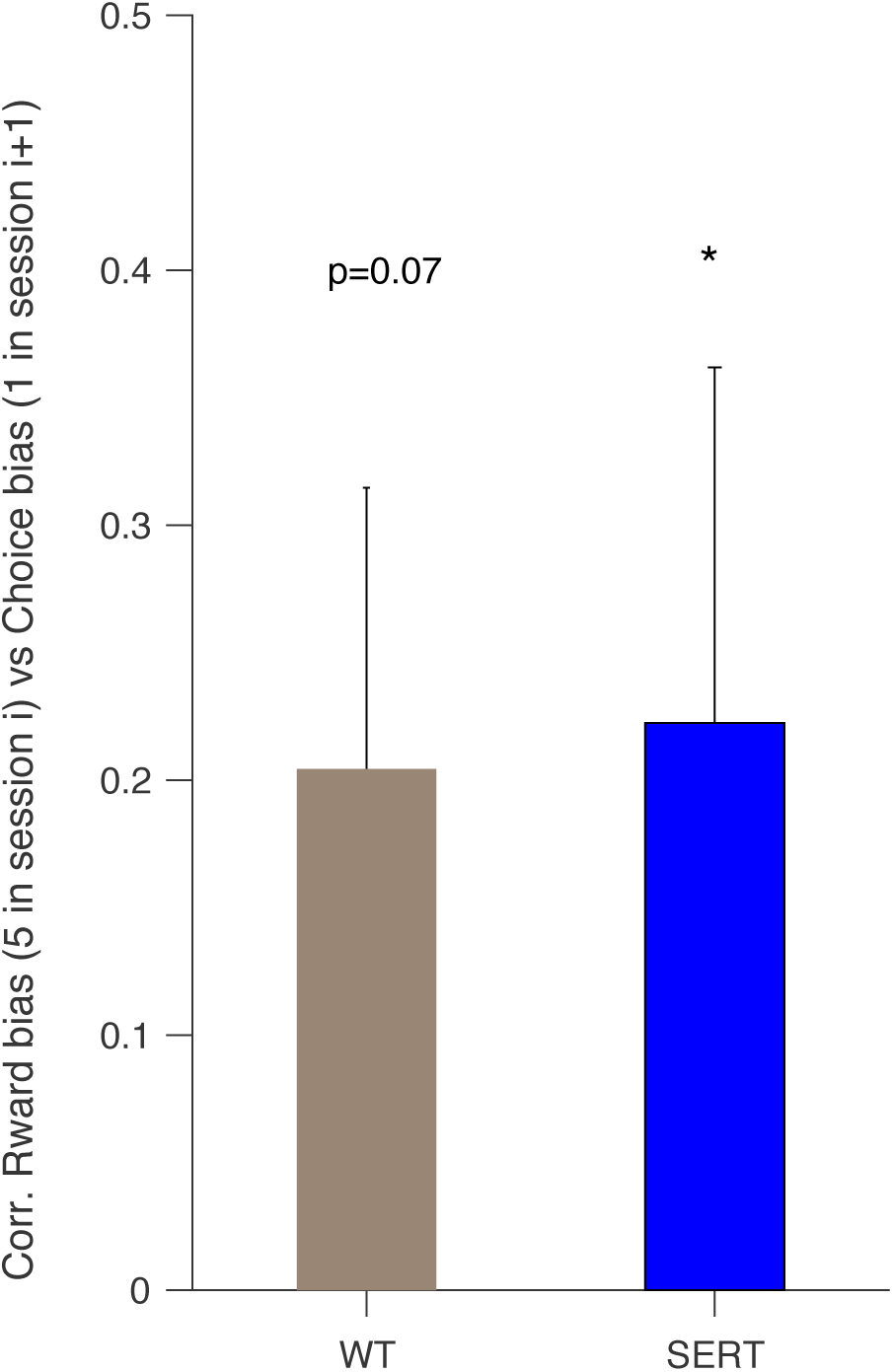
Choices following long ITIs in the first quintile were correlated with reward bias in the fifth quintile in the previous session. The star indicate how significantly the correlation is different from zero, tested by a permutation test. The test statistic was constructed by the mean of the correlation coefficients of four animals, where the correlation coefficient was computed by randomly permuted data in each animal. One star indicates *p* < 0.05. The error bars indicate the mean ± SEM of data.

